# Anxiety, avoidance, and sequential evaluation

**DOI:** 10.1101/724492

**Authors:** Samuel Zorowitz, Ida Momennejad, Nathaniel D. Daw

**Affiliations:** Princeton Neuroscience Institute, Princeton University, Princeton, NJ 08540; Department of Biomedical Engineering, Columbia University, New York, NY 10027; Princeton Neuroscience Institute, and Department of Psychology, Princeton University, Princeton, NJ 08540

**Keywords:** anxiety, avoidance, fear generalization, decision theory, computational psychiatry

## Abstract

Anxiety disorders are characterized by a range of aberrations in the processing of and response to threat, but there is little clarity what core pathogenesis might underlie these symptoms. Here we propose that a particular set of unrealistically pessimistic assumptions can distort an agent’s behavior and underlie a host of seemingly disparate anxiety symptoms. We formalize this hypothesis in a decision theoretic analysis of maladaptive avoidance and a reinforcement learning model, which shows how a localized bias in beliefs can formally explain a range of phenomena related to anxiety. The core observation, implicit in standard decision theoretic accounts of sequential evaluation, is that the potential for avoidance should be protective: if danger can be avoided later, it poses less threat now. We show how a violation of this assumption — via a pessimistic, false belief that later avoidance will be unsuccessful — leads to a characteristic, excessive propagation of fear and avoidance to situations far antecedent of threat. This single deviation can explain a range of features of anxious behavior, including exaggerated threat appraisals, fear generalization, and persistent avoidance. Simulations of the model reproduce laboratory demonstrations of abnormal decision making in anxiety, including in situations of approach-avoid conflict and planning to avoid losses. The model also ties together a number of other seemingly disjoint phenomena in anxious disorders. For instance, learning under the pessimistic bias captures a hypothesis about the role of anxiety in the later development of depression. The bias itself offers a new formalization of classic insights from the psychiatric literature about the central role of maladaptive beliefs about control and self-efficacy in anxiety. This perspective also extends previous computational accounts of beliefs about control in mood disorders, which neglected the sequential aspects of choice.

## 1 Introduction

Though anxiety disorders differ in their particular symptomology, and in the content and situations that elicit symptoms, they all are similarly characterized by aberrations in the processing of and response to threat^1^. In particular, at least three symptoms manifest across many of the anxiety disorders. First, anxiety is associated with exaggerated threat appraisal, or a bias towards evaluating threat as disproportionately greater in likelihood and severity than is warranted^2^. Second, anxiety is also associated with fear generalization, wherein the primary threat becomes associated with increasingly distal locations, events, and thoughts^3^. Finally, anxiety is associated with persistent avoidance behavior, which often occurs well in advance of the materialization of actual threat^4^. (Here we distinguish between avoidance and escape, where the former describes actions taken to prevent the onset of threats whereas the latter describes defensive responses to proximal threat.) Excessive avoidance behaviors are an especially harmful aspect of anxiety disorders, both because they interfere with daily life and because they indirectly maintain anxiety by preventing learning from the non-occurrence of perceived threats. Though laboratory studies of decision making and learning in anxious populations have corroborated these clinical observations^5,6,7^, none offer an explanation as to the root of these symptoms.

These symptoms are particularly puzzling from a decision theoretic perspective^8^. In many circumstances, distant threat should not impinge upon decision making in the present. Indeed, we argue that fear and avoidance of situations far in the future violates the basic logic of evaluation over sequential trajectories of action. This is because avoidance is by nature protective: the ability successfully to avoid danger in the future means an agent need not also do so now. For instance, cars endanger pedestrians but can be reliably avoided by following traffic signals; given that, staying indoors offers little or no additional protection from accidents. This is an instance of a fundamental property of evaluation in sequential decision making: the value of present action turns fundamentally on assumptions about subsequent events, which importantly include the agent’s own subsequent choices. Typically, it is appropriate to assume that an agent will continue to make good (i.e. reward-maximizing/harm-minimizing) choices down the line, and that good choices at the current stage should therefore anticipate this.

This line of reasoning hints that a fundamental aberration in anxiety disorders may relate to this assumption, which otherwise should preclude the spread of threat to antecedent situations and subsequent excessive avoidance. Indeed, anxiety disorders are associated with pessimistic beliefs about the future^2^. Clinically and subclinically anxious individuals judge future threat as more likely than do non-anxious individuals^9,10,11^. Importantly, the development and maintenance of clinical anxiety is strongly tied to diminished perceived control^12,13,14^, such that anxious individuals are more likely to endorse the belief that they are unlikely or unable to mitigate future threat. Indeed, lack of belief in one’s ability to successfully navigate future danger is associated with anxiety^15,16^, and an increased belief in perceived control over threat is correlated with symptom reduction across the family of anxiety disorders^17^.

Here, we develop this idea, that symptoms of anxiety may arise from misbeliefs about future avoidance, into a formal model of evaluation under pessimistic assumptions about future choices. We show that a single, localized deviation from normative evaluation can explain a surprising range of features of anxious behavior, including exaggerated threat appraisal, fear generalization, and persistent avoidance. This account also offers a new formalization of classic insights from the psychiatric literature about the central role of beliefs about control and self-efficacy in anxiety^12,13^. Specifically, we show through simulation that a model with a misbelief about the reliability of future self-action gives rise to a number of characteristic symptoms and laboratory results concerning anxiety. Our perspective also extends previous computational accounts of beliefs about control in mood disorders (e.g.^18^), which neglected the sequential aspects of choice.

## 2 Model description

We model anxious decision making in the context of Markov decision processes (MDPs). A standard normative assumption is that agents attempt to optimize the expected cumulative discounted reward:

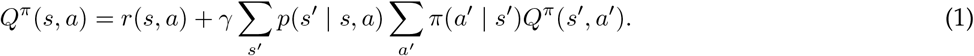

For any particular state-action (*s, a*), this is necessarily defined relative to a policy *π*(*a*′ | *s*′) specifying the assumed distribution of *future* choices. (It is also defined relative to a discount rate *γ*, controlling the present value of future outcomes.) The return can be optimized self-consistently under the assumption that the agent makes the return-maximizing choice at each step in the future, leading to the familiar expression for the optimal values,

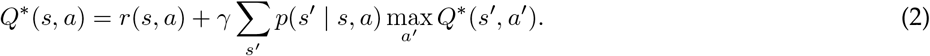

The “max” operator in Eq. 2 yields a fundamental asymmetry between approach and avoidance, as illustrated in Fig. 1. It formalizes the assumption that the agent makes the return-maximizing choice at each step. Through this operation, opportunities for reward (which maximize the argument) propagate recursively to earlier steps, but avoidable dangers do not, because the return-maximizing action is to avoid. (To the extent obtaining reward or avoiding harm are only imperfectly achievable, this propagation and attenuation at each step are graded according to success probability, but the basic asymmetry remains.) This principle is highlighted in a toy MDP (Fig. 1): a deterministic open gridworld with two terminal states, a rewarding state and a punishing one (Fig. 1a). The optimal state values *V*\* = max_*a*_ *Q*\*(*s, a*) (Fig. 1b) reflect a “mountain” of opportunity propagating recursively from the reward. Conversely, because harm is avoidable in this environment, its negative value is contained: all states (even those adjacent to threat) represent the positive opportunity for reward.

**Figure 1:**
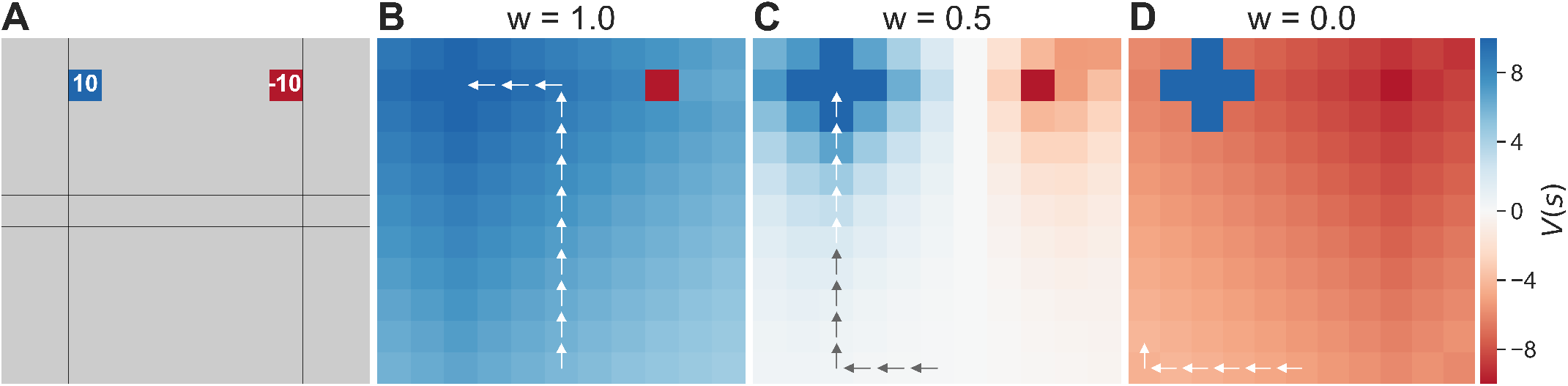
(A) A simple deterministic gridworld with two terminal states: one rewarding (blue) and one aversive (red). (B-D) States colored by their value under different levels of pessimism, with arrows showing an optimal trajectory. (B) For an optimistic agent (*w* = 1), all states (other than the harmful state) take on positive value. (C) For a pessimistic agent (*w* = 0.5), negative value spreads from the source to antecedent states. (D) With increasing pessimism (*w* = 0), the extent of the spread grows worse, and the return-optimizing trajectory becomes more distorted and avoidant. (Parameters: *γ* = 0.95)

Although the return-maximizing assumption self-consistently defines optimal behavior, an agent need not be restricted to it^19^, and might in principle anticipate encountering danger under different (e.g., pathological) assumptions about the future. For example, an agent may expect to fail to take the correct protective actions in later states (i.e., to use a sub-optimal *π*(*a* | *s*)), or may believe the world’s future dynamics do not guarantee reliable avoidance even so (i.e., under stochastic or adversarial transition dynamics *P*(*s*′ | *s, a*)). Consider an agent who has such pessimistic expectations about dangerous events at future steps. Note that assumptions of this sort, even if incorrect, can serve adaptive purposes. In general, pessimistic assumptions can help to ensure robustness and safety under uncertain or even adversarial scenarios^20^. Related work in RL shows how computing returns with pessimistic predictions can help to quantify variability in outcomes (i.e., to learn different points in the distribution of possible returns)^21^, which is one way to explicitly and tunably take account of risk tolerance. Indeed, it is common in machine learning theory to optimize outcomes under worst-case assumptions.

Here we propose unrealistically pessimistic assumptions as a root cause of many anxious symptoms. Such pessimism can be encoded either in the policy, *π*(*a* | *s*), or transition probabilities, *P*(*s*′ | *s, a*). These, respectively, correspond to misbeliefs about either one’s own avoidance actions or the environment’s responses to them, a point we return to in discussion. Here for concreteness we focus on distortions in the policy. In particular, we adopt the *β*-pessimistic value function from^22^, to define state-action value in expectation over a mixture of the best and worst action:

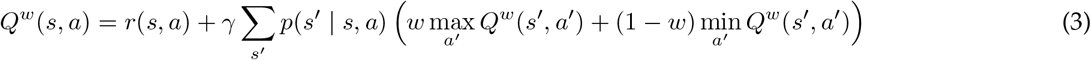

The weight parameter *w* controls the degree of pessimism. An optimistic agent (*w* = 1) expects in the future to act fully in accordance with its preferences, whereas a pessimistic agent (*w* = 0) expects to act contrary to its preferences. Importantly, this belief is false under the model: that is, we assume, at each step, that the agent actually chooses by maximizing the values given by Equation 3. But these values are always computed under the assumption that the agent will then not maximize at all later steps. (Another way to say this is that we assume choices at each step are optimal, but only under the assumption that later choices will not be.) Fig. 1c,d illustrates the the consequences of different levels of pessimism for valuation in an example gridworld. The value of threat propagates, with increasing distance across the state space, due to an increasing expectation that the agent may fail to avoid threat in the future.

This simple simulation reflects a localized violation of the core decision theoretic assumption of future return-optimizing action. Therefore, the model’s behavior already echoes several core symptoms of anxiety disorders. Namely, the pessimistic agents in Fig 1c,d exhibit exaggerated threat appraisals (otherwise neutral states unrealistically signal danger); generalization of fear (threat value spreads across the gridworld); and persistent avoidance (early on, the agent takes paths which maintain increasing distances from threat). Importantly, as we will elaborate below, this deviation from the usual assumptions is supported by prominent clinical theories of anxiety.

## 3 Simulations

In what follows, we will demonstrate through simulation how our simple model can account for anxious behavior in laboratory-based studies of *sequential learning* and decision making. (Because in our model anxiety arises through biased sequential evaluation, we will not address one-step bandit tasks where others have reported learning deficits associated with anxiety^5,7^.) We will also show that our model is consistent with clinical theory describing the transition from clinical anxiety to depression. Unless otherwise noted, state and action values under varying degrees of pessimism were solved for using the value iteration dynamic programming method^23^. All simulations were implemented in the Python programming language and the code is publicly available at https://github.com/ndawlab/seqanx.

### 3.1 Approach-avoidance conflict

One behavioral finding characteristic of anxiety disorders is unbalanced processing of approach-avoidance conflict^27^. Anxious individuals are more likely to forgo potential gains in order to avoid potential danger. Many of the disruptions anxiety causes to everyday functioning (e.g. avoiding social obligations for fear of possible social humiliation) can be understood in these terms. As such, many have sought to probe and measure this behavior in the laboratory. For instance, in the balloon analog risk task (BART)^24^, participants attempt to earn money by pumping virtual balloons. With each pump, the balloon inflates and money is earned, but so too does the chance that the balloon pops and the accumulated earnings are lost. At any point in a trial, a participant may cash out, banking the money earned and ending the trial. Anxiety is correlated with fewer pumps of the balloon and earlier cash-outs in the BART^25,26^.

As shown in Fig. 2, our model easily accommodates this result. Whereas optimistic agents pump until the marginal gain of a pump no longer offsets the chance of the balloon bursting, optimal choice under increasingly pessimistic (i.e., anxious) assumptions cashes out progressively earlier – similar to empirical findings^25,26^. This is because it anticipates and avoids future errors in choice, which would otherwise result in the balloon popping. Our model can analogously explain other manifestations of biased approach-avoid conflict in anxiety, such as in the predator avoidance task^28^.

**Figure 2:**
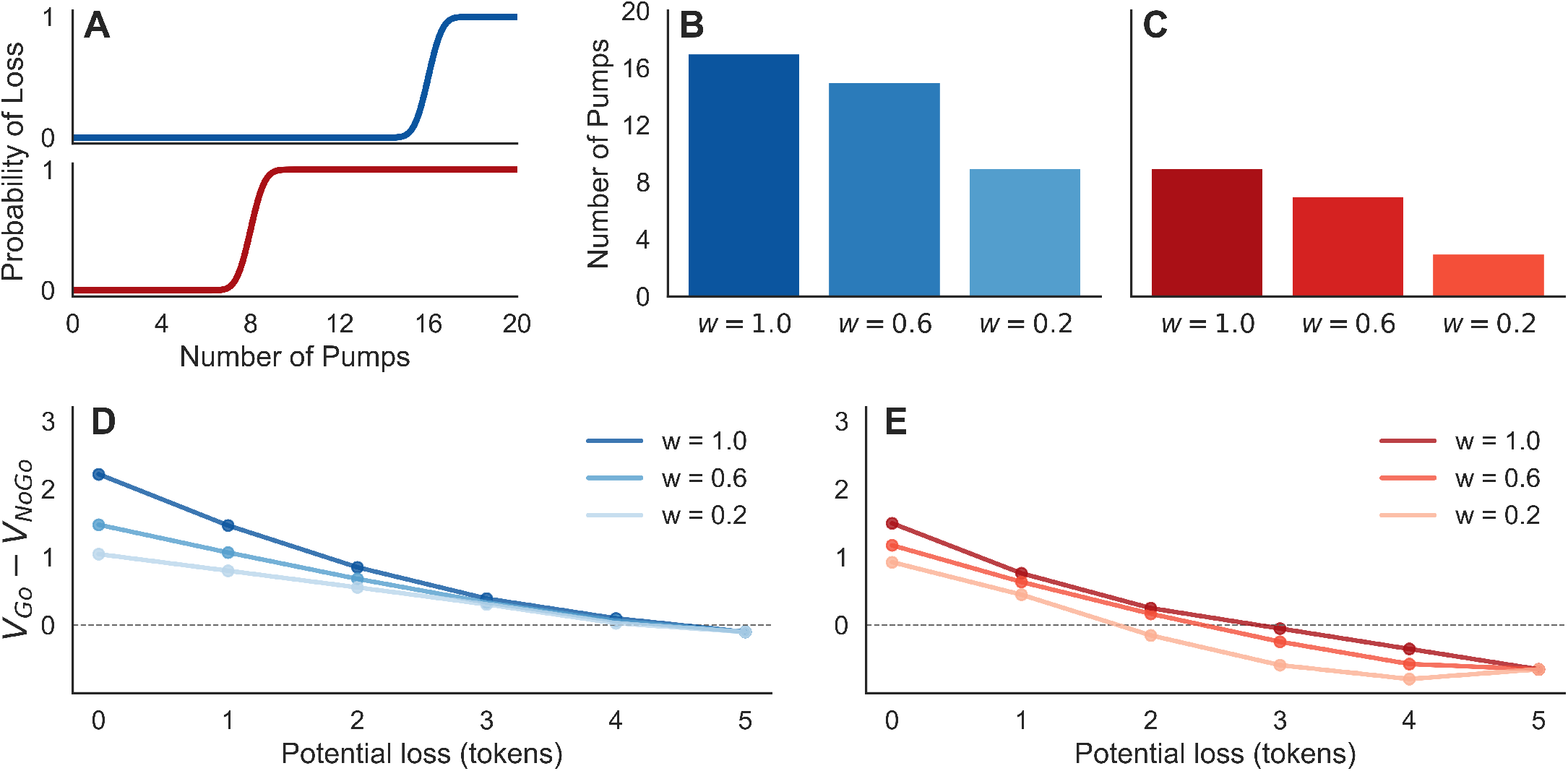
(A) The Balloon Analog Risk Task (BART)^24,25,26^. The risk of balloon burst (point loss) increases with each pump and does so earlier for the high risk (red) than low risk (blue) balloons. (For full rules of the task, see Methods.) The optimal policy (number of pumps) under increasingly pessimistic valuation is presented for low (B) and high risk (C) balloons. The optimistic agent (*w* = 1) prefers a policy reflecting the true environmental risk. The moderate (*w* = 0.6) and strongly (*w* = 0.2) pessimistic agents cash-out earlier, as is observed in anxious individuals. (D,E) In the sleeping predator task, the risk of loss is constant but the cost of loss still increases as more rewards are gathered. The value of reward pursuit under increasingly pessimistic valuation is presented for scenarios with low (D) and high risk (E) of predator awakening. The relative value of approach (vs. avoid) decreases with loss amount and threat level, and moreso under pessimistic assumptions. (Parameters: *γ* =1.0)

A unique prediction of the model (because we assume optimal choice under the assumption of *future* suboptimality) is that bias should arise only when beliefs about *future* avoidance are involved, rather than direct conflict between immediate impulses. Recent data^28^ using a predator avoidance task, analogous to the BART, support this view. In this task, increasing trait anxiety predicted earlier escape (analogous to cash-out in the BART) for slow predators (for whom future decisions to escape were a relevant consideration) but not for fast ones (who would attack immediately, mooting consideration of future steps).

Relatedly, the model can also capture findings of increased behavioral inhibition, measured as prolonged response times, in anxious individuals under threat^29^. In the behavioral inhibition task, participants seek tokens adjacent to a virtual “sleeping predator”, which are all lost if the predator awakes. Though in this variant the risk of predation is constant throughout a trial (rather than increasing, as in the BART) the potential loss from capture still increases with each token collected. Bach^29^ finds that participants are slower to collect tokens as this potential loss increases, and that this slowing is enhanced by subclinical anxiety. We can capture this effect in our model by noting that the relative value of approach compared to avoidance is reduced as potential loss and the risk that the predator awakes increase (Fig. 2d/e). These effects are amplified under more pessimistic (i.e., in our model, more anxious) assumptions about future actions. Thus, as before, anxious pessimism in our model produces greater and earlier choice of avoidance, analogous to earlier cashout in the BART. To extend this effect to reaction times, we adopt the further, standard assumption that actions (here, approach) are slower when their relative value compared to alternatives is lower^30^. In this case, the model captures behavioral inhibition (slower responses as threat increases), and its enhancement by anxiety, as measured by Bach. (Note that we need not assume the coupling of reaction times to action value spread is due to difficulty in decision formation per se, which Bach argues against: it may, for instance, reflect Pavlovian initiation biases^31^.)

### 3.2 Aversive pruning

Another laboratory phenomenon associated with anxiety is “aversive pruning” in planning^32,33^. This refers to the idea that when evaluating future action trajectories in a sequential task like chess, people are resource-limited, cannot evaluate all possible options, and must selectively consider certain paths and neglect others. One proposal for how people accomplish this is aversive pruning^32^, wherein choice sequences involving large losses are discarded from further evaluation. An example of aversive pruning is shown in Fig. 3a. Although the optimal choice in the decision tree from is to weather an initial large loss (e.g., −70) in order to reap the large gain that follows, people tend to disfavor this path suggesting they prune it and consequently neglect the later gain. The degree of such pruning correlates, depending on the study, with subclinical depressive^32^ or anxiety^33^ symptoms.

**Figure 3:**
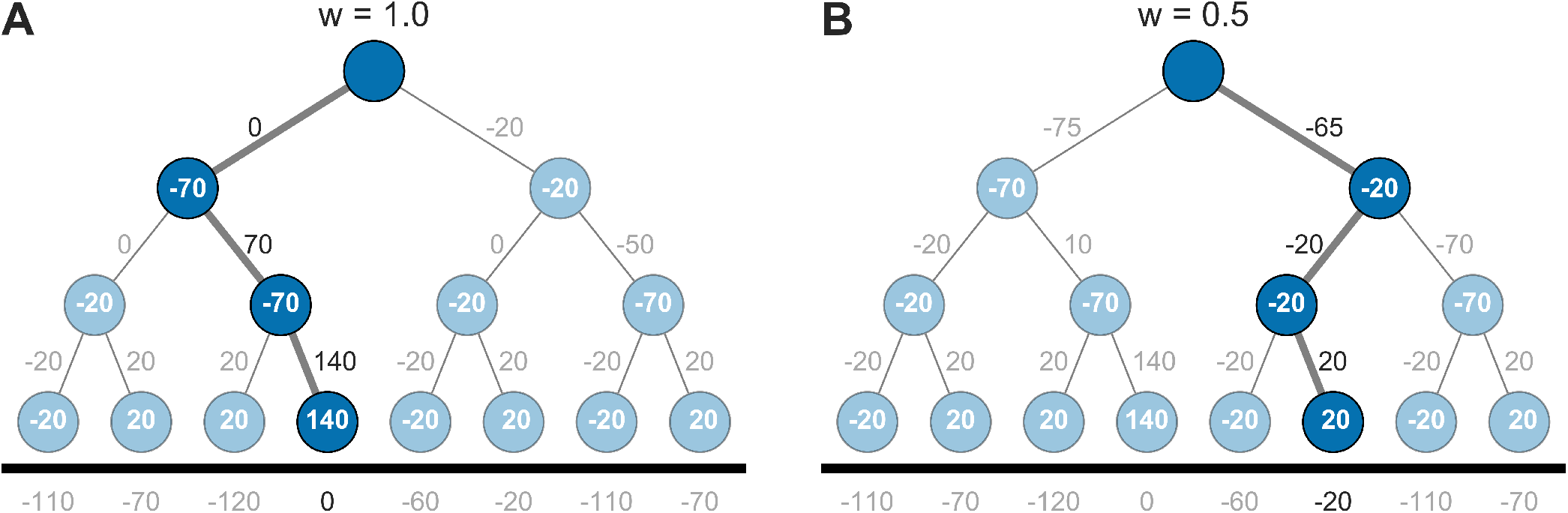
(A) The decision tree environment^32,33^. An optimistic agent (*w* = 1) prefers the optimal loss-minimizing policy through the initial large loss (left branch). (B) A pessimistic agent (*w* = 0.5) comes to prefer the branch without the large loss so as to avoid being unable to recoup the large initial loss. One-step rewards (or costs) are presented in each state; the net value *Q* on each path is shown numerically. (Parameters: *γ* = 1.0)

Our model predicts this result (Fig. 3b) as specifically linked to our model of anxious pessimism, though for a somewhat different reason than in Huys’ original modeling. In our model, pessimistic (anxious) agents neglect large gains deeper in the tree, not because they fail to consider them (here we assume full evaluation of the Bellman equation), but because with increasing anxiety they increasingly expect the potential of choosing incorrectly afterward, thus failing to recoup the loss (and, mathematically, probabilistically pruning the better branches). Future research could use slight variants in the decision trees to tease apart these different interpretations, for example, by comparing decision trees that differ only in what follows the large initial loss. Under a model of aversive pruning, such a change should not impact the proportion of agents selecting the left branch; in contrast, our model predicts choice should parametrically increase with the extent of the amelioration.

### 3.3 The anxiety-depression transition

So far, we have considered only the asymptotic preferences implied by our pessimistic value function, which we computed directly through value iteration. But we can also consider the process of learning under this value function (e.g., by variants of Q-learning^23^ or DYNA^34,35^ using the *β*-pessimistic return). The dynamics of such learning may speak to the progression of symptoms.

Of note in this respect, anxiety and depression are highly comorbid, with almost half of individuals with a lifetime depression diagnosis also diagnosed with an anxiety disorder^36^. One notable proposal is that this association often (though by no means exclusively) arises longitudinally: in particular, that clinical anxiety precedes certain types of depression^37,38^. The idea, in brief, is that uncertainty in one’s ability in the face of future threat results in anxiety and avoidance behaviors. Persistent avoidance, in turn, begets foregone reward, and ultimately to a belief that reward is unobtainable and subsequently depression. This informal story can be captured by simulations of learning in our model (Fig. 4) in environments like that of Fig. 1. Over the course of learning, the penumbra of negative value under pessimistic assumptions spreads gradually throughout the environment. This can in turn lead the agent to expect no reward and, also echoing the anergic symptoms of depression, forego action altogether. This last point in particular dovetails nicely with theoretical accounts of the anergic aspects of depression^8^, which point out that low experienced reward rates should in decision theoretic accounts lead to reduced response vigor^31^ leading to a potentially self-reinforcing downward spiral.

**Figure 4:**
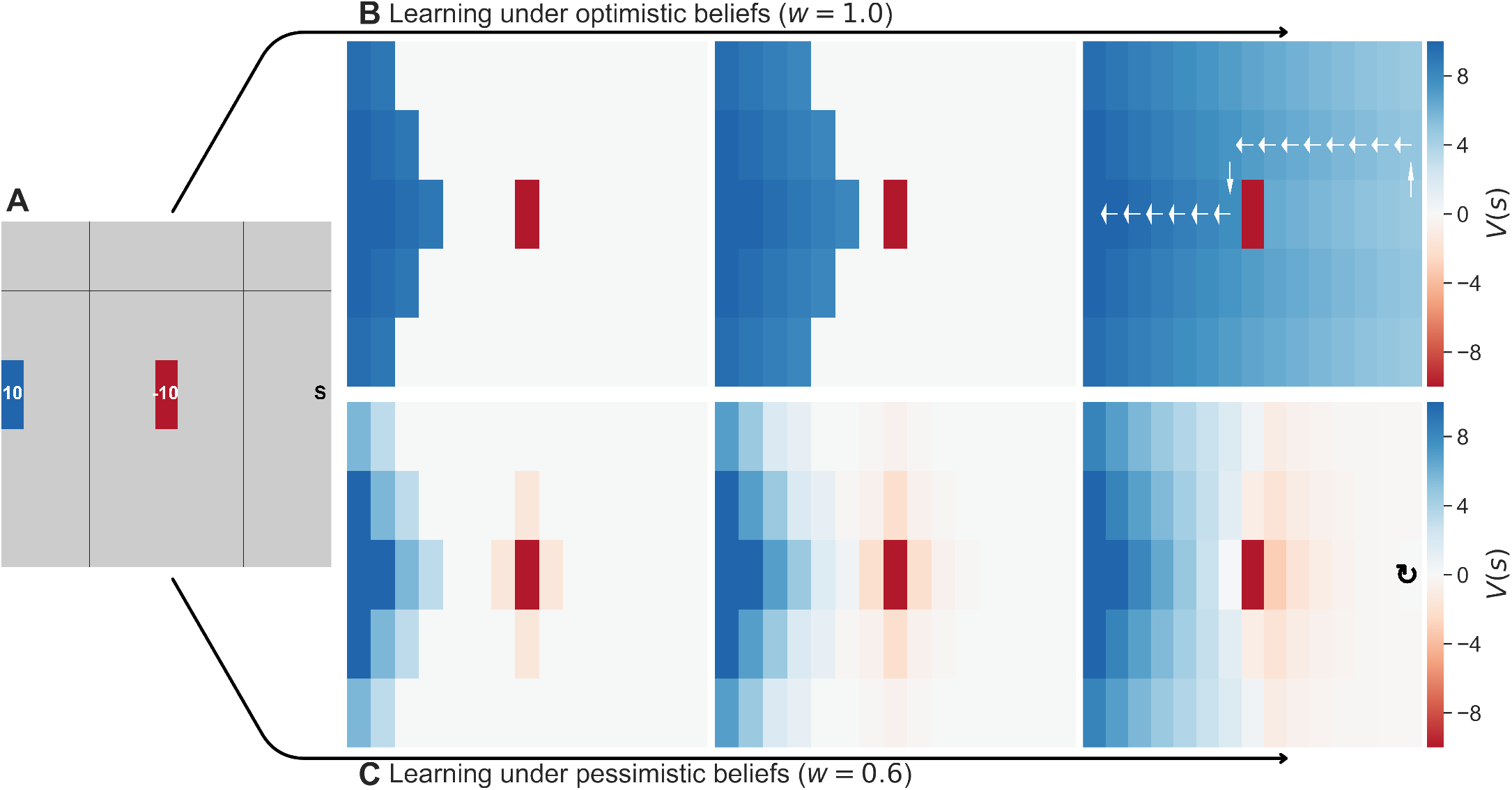
(A) A simple deterministic gridworld with two terminal states: one rewarding (blue) and one aversive (red). (B, c) The development of value expectancies over three steps of learning, for two levels of pessimism. States are colored by their value under different levels of pessimism, with arrows showing an optimal trajectory. (B) For an optimistic agent (*w* = 1), all states (other than the harmful state) take on positive value with learning. (C) For a pessimistic agent (*w* = 0.6), negative value spreads from the source to antecedent states. As a result of avoidance, the agent learns reward is unobtainable and develops anergic symptoms (i.e. foregoes action). (Parameters: *γ* = 0.95)

In addition to suggesting one explanation for the comorbidity of anxiety and depression, our model also hints at a reason for the longevity and recurrence of anxiety disorders even with treatment. Because pessimistic expectations allow for threat value to spread to states and actions far antecedent of the primary danger (e.g., Fig. 1d), it would accordingly also take a great many steps of iterative learning to correct all these exaggerated appraisals of threat. Frustratingly, these biased estimates of value may still remain even after a misbelief in the efficacy of future action is corrected for in a course of therapy. This phenomenon (similar to failures of model-free RL algorithms to adjust to reward revaluation without extensive relearning;^39^) may offer at least a partial answer to a classic puzzle in pathological avoidance, i.e. why it is so resistant to extinction^40^, and to the unfortunately high rates of anxiety recurrence following treatment^41^.

### 3.4 Free choice premium

Finally, the model also offers a novel prediction tying anxious beliefs to a classic, but hitherto separate, phenomenon known as the free choice premium. This refers to the finding that, all else being equal, people tend to treat choice as itself valuable: i.e., choices which lead to more choice opportunities in the future are preferred to those that lead to fewer future choice opportunities. A free choice premium has been observed in multiple behavioral experiments^42,43^. A variant of a free choice premium paradigm from two previous studies^44,45^ is presented in Fig. 5a. In the task, participants repeatedly choose between a free choice option, allowing for an additional future choice, and a fixed choice option. Importantly, both choices lead to identical, stochastic outcomes (e.g. 50/50 chance of [1, −1]). Empirical studies have found that human subjects (from a general, healthy population) prefer the free choice option despite it conferring no additional benefits relative to the fixed choice option.

**Figure 5:**
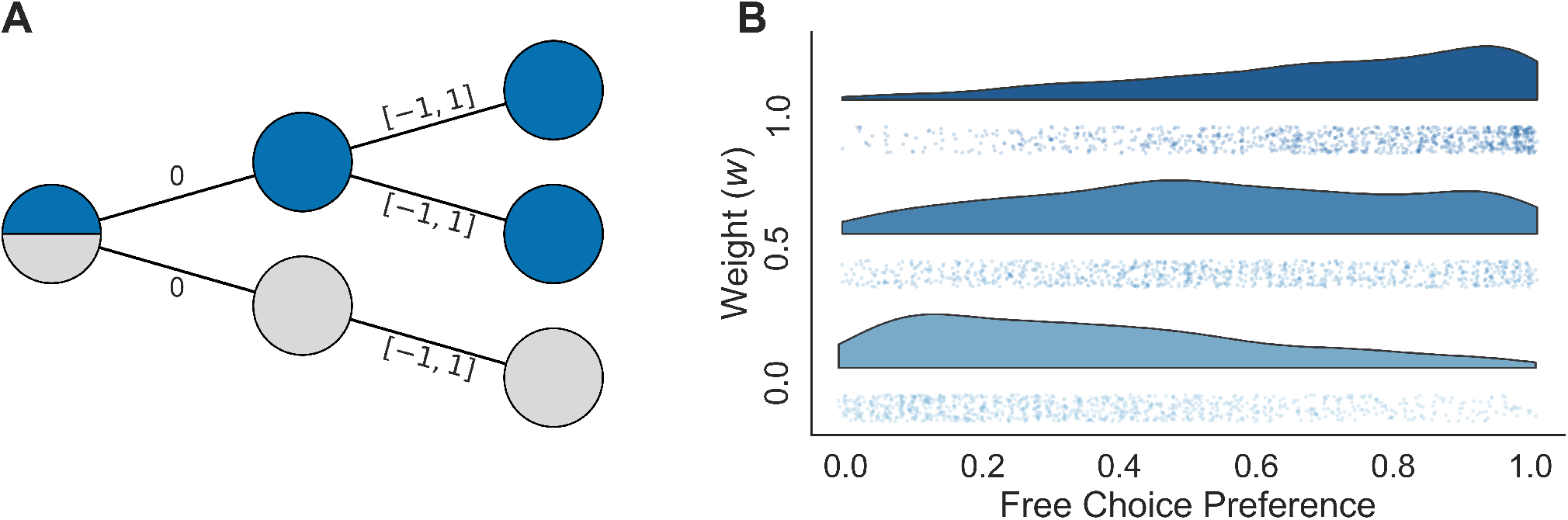
(A) The free choice premium task^44,45^ with equal chance of outcome *R* ∈ [1, −1]. Non-anxious participants exhibit a preference for the free choice option (blue) despite it conferring no benefit over the fixed choice option (grey). (B) Pessimistic agents show an attenuated free choice bias. The fractional preference for the free choice option over the simulated experiment is shown for three populations of subjects with levels of pessimism *w* (y-axis; each dot represents an individual simulated agent, and the smoothed density of free choice bias for each pessimism level is also shown.) (Parameters: Q-learning with *γ* = 1.0, and inverse temperature, *β*, increased from 1 to 15 over 100 episodes.)

On one account^43^, this free choice preference directly and specifically reflects the assumption about sequential choice whose violation we argue is core to anxiety: i.e. that the agent will make reward-maximizing choices in the future. Under such an optimistic assumption (and given noisy and imperfect knowledge about the value of each option, due to learning), additional options are valuable in the sense that free choice can be expected to exploit the best among them. Namely, the maximum over several noisy values is, in general, larger than a single option from the same distribution. Our proposal makes the novel prediction that, if as we hypothesize anxiety reflects a violation of this optimistic assumption, then anxious individuals will exhibit a diminished or reversed free choice bias, as shown in simulation in Fig. 5b. Future empirical research will be required to test this prediction.

## 4 Discussion

Central to anxiety disorders are symptoms including exaggerated threat appraisal, threat generalization, and excessive avoidance^2,3,4^. We have presented a simple computational account suggesting how a single underlying pessimistic misbelief can give rise to these aberrations in learning and choice. We use a reinforcement learning approach in which undue pessimism results in maladaptive policy. Specifically, we showed how a failure to believe in the reliability of one’s future actions can effectively backpropagate negative value across states of the environment. This process results in a range of inferences and behaviors resembling those observed in clinical anxiety. Though it is by no means a complete account of anxiety, our account ties together a surprisingly wide range of symptoms of anxiety disorders.

We are not the first to propose a formal theory of control in psychiatry using MDPs. Huys & Dayan^18^ also provided a computational account of learned helplessness through simple models of one-step action-outcome contingencies. Our accounts differ particularly in our exclusive focus on control in the sequential setting, which Huys & Dayan did not address. Indeed, we propose that ultimately key to anxiety is precisely the way in which evaluation in sequential tasks is necessarily reliant on expectations about future choice and events. Similarly, research and modeling by Bishop and colleagues^46,47^ also taking a decision theoretic approach has stressed the importance of uncertainty as a core feature of anxiety. Specifically, they have described uncertainty as inherently aversive in anxiety, and have presented models of how uncertainty may be increased in anxiety (e.g. aberrant processing of environmental volatility). The present work is compatible with deficits in processing uncertainty, and might instead be viewed as an attempt to unpack why uncertainty is aversive: because, in our view, it is resolved (i.e. marginalized) under pessimistic distributional assumptions. As for control, we extend this view to focus on how uncertainty is resolved in the sequential setting, and also to zero in on particular instances of uncertainty (about future actions, and some other options discussed next) and misbeliefs about them that, we argue, are particularly consequential.

For concreteness, we formalized pessimistic assumptions in terms of only one of several variants of a more general family of models, but we do not mean this restriction as a substantive claim. In particular, we focused on the agent’s beliefs about their own future actions: expecting failure in handling or avoiding future threat. However, this is just one of several different pessimistic misbeliefs that could satisfy the basic logic of our model and produce similar symptoms. These other beliefs need not be mutually exclusive, though they might reflect different cognitive routes to symptoms that would, in turn, imply different psychotherapeutic strategies.

For instance, one variant of the model is suggested by the observation that Eq. 1 is also computed in expectation over the anticipated future environmental dynamics *p*(*s*′ | *s, a*). Thus, pessimism can alternatively be encoded in this distribution: e.g., a false belief that the world’s response to one’s choices is unpredictable or adversarial. Because the Bellman equation for the return averages over this distribution in addition to the choice policy at each step, and because an unpredictable environment also reduces the efficacy of avoidance, either formulation can produce ultimately similar results in our simulations here. A third variant of our model arises from uncertainty about the current state s of the environment. Although we have taken it as fully observed, if the world state is only partly known, then this distribution too must be averaged out in evaluating each action^48^, and here also a pessimistic skew will propagate the expectation of danger and result in exaggerated avoidance^49^. In summary, pessimistic resolution of several different varieties of uncertainty (e.g., about future action, environmental dynamics, or environmental state) could each produce similar symptoms for analogous reasons. However, from the perspective of cognitive theories of anxiety, these represent quite different maladaptive beliefs: a key difference that may be relevant in guiding treatment (especially cognitive psychotherapies aimed at ameliorating the false beliefs) of a host of anxiety disorders.

Particularly due to the way it encompasses several such variants, our account formalizes a longstanding range of theory on the role of control in anxiety. Central to many prominent cognitive theories of anxiety in the psychiatric literature is a perceived lack of control. For example, self-efficacy theory^12^ and the triple vulnerabilities model^13^ both posit that a reduced belief in the ability to effectively respond to future threat is involved in the genesis and maintenance of clinical anxiety. In contrast, and focused less on the self, the learned helplessness theories of anxiety^37^ claims clinical anxiety results from an uncertain belief in the controllability of the environment, such that future threat cannot be effectively mitigated or avoided. As we note above, the present model and analysis (though simulated above in terms of selfefficacy) can accommodate either variant, and shows how they relate to one another as members of a more general family of accounts.

The possibility of multiple anxious phenotypes, each characterized by unique but not mutually exclusive beliefs, suggests the need for behavioral assays designed to isolate and interrogate such biases. One such task is the free choice premium paradigm described above, which captures pessimism (or optimism) about one’s own choices. An analogous task might measure pessimistic expectations about environmental state transition probabilities. For instance, this could be accomplished in a variant of sequential decision making tasks that require subjects to learn the transition structure of a multi-step decision tree^50,51^ and make choices to gather rewards in it. Pessimistic expectations about environmental state transitions would bias choices in this type of task. Individually, these tasks could test our hypothesis that either sort of misbelief is associated with symptoms of anxiety; they could also be compared to one another (and to more detailed self-report assessments of beliefs about control or self-efficacy) to investigate potential heterogeneity across patients in the antecedent of anxiety.

Importantly, although we have considered the most pathological cases of pessimism, these may reflect the exaggerative extremes of an otherwise adaptive evaluation strategy. Traditionally, the goal of reinforcement learning algorithms is to a find reward-maximizing policy with respect to the expectation (average) of returns. However, depending on one’s risk attitude and uncertainty about the environment (e.g. if there is potential for catastrophic loss), it may be instead preferable to learn a policy with respect to an alternative and more pessimistic objective function, similar to the one considered here. Accordingly, returns and agent behaviors similar to ours arise in previous research on learning risk-sensitive and robust policies^52,53,21^.

We have centered our discussion at Marr’s^54^ computational level: on beliefs and their consequences in terms of action values. We have so far remained agnostic as to how, algorithmically or mechanistically, these misbeliefs are implemented in the brain.^55^ Importantly, the brain is believed to contain multiple distinct mechanisms for evaluating actions (e.g, model-based and model-free learning;^39,8^), and pessimistic beliefs might play out either differentially or similarly through each of these mechanisms. One promising possibility is that these symptoms mainly reflect aberrations in model-based planning^8^, i.e. explicitly evaluating actions by mentally simulating possible trajectories. Recent work suggests this process may be accomplished by mentally “replaying” individual potential trajectories^56,57^. In this setting, the biases we suggest would amount to over-contemplating, or over-weighting, certain pessimistic trajectories^58^. Such a bias might be detectable using neuroimaging, as a change in which types of events that tend to be replayed^56,59^. Such a biased replay process, in turn, may also correspond to worry and rumination. Indeed, in line with the present results, chronic worry is associated with reduced perceived control, diminished belief in self-efficacy in response to threat, and exaggerated threat appraisal^60^. This suggests that clinical anxiety may in part result from planning processes gone awry.

It is important to note that the present model may not describe all anxiety disorders with equal accuracy. Indeed, our analysis of pessimistic sequential evaluation is, by definition, a model of prospective cognition. Thus, the present results are more likely to accurately describe the anxiety disorders which primarily involve aberrations in future-oriented cognitive processes, such as generalized and social anxiety disorders. Naturally, the present model can account neither for the compulsive behaviors of obsessive-compulsive disorder (OCD) nor the memory disturbances of post-traumatic stress disorder (PTSD). That said, recent clinical studies suggest that diminished perceived control is a vulnerability factor common to all anxiety disorders, including OCD and PTSD^14,17^. Importantly, aspects of other psychiatric disorders that involve future-oriented misbeliefs, worry, and avoidance behaviors (e.g. eating disorders^61^) may similarly be well-described by the current account. Indeed, and much more speculatively, the model may also have implications for bipolar disorders. The onset of manic symptoms is associated with overoptimistic perceived control^62^, and our same model and basic reasoning (though now envisioning excessively optimistic rather than excessively pessimistic beliefs) may help to explain how this bias may translate into risk-seeking behavior and dysreguated goal pursuit. As such, the present results are transdiagnostic and not limited to one particular diagnosis.

Finally, the relationship between evaluation, planning, and neural replay discussed above suggests potential future work that might help to bring this model into contact with the more memory-related aspects of PTSD. For instance, the same replay processes that can be used to evaluate actions can also, in theory, update predictive representations, cognitive maps, or models of the environment^35^, such as the successor representation^63,64^. If so, similar biases in replay could result in not just aberrant avoidance behavior, but also progressive, aberrant remodeling of world models or cognitive maps, an observation which may connect to the rich and complex set of issues on memory involvement in PTSD.

## 5 Methods

We model anxious decision making in the context of Markov decision processes (MDPs). Tasks were modeled as deterministic, infinite horizon, discrete time environments. Some (detailed below) were modeled with discounted returns *γ* < 1. All simulations were implemented in the Python programming language and the code is publicly available at https://github.com/ndawlab/seqanx.

For all but the free choice premium task, we defined state-action values, *Q*(*s*, *a*), in accordance with our modified, pessimistic Bellman equation Eq. 3, reproduced below for convenience:

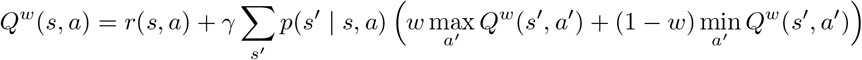

Here, *Q*-values were solved for directly through value iteration^23^. By contrast, *Q*-values in the free choice task were computed using the *β*-pessimism temporal-difference learning algorithm^22^, where the reward prediction error is defined as:

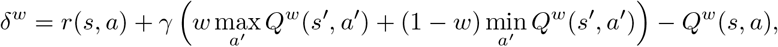

and the update rule is defined as:

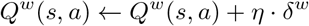

where *η* is a learning rate. The parameterizations of the MDP environments and learning algorithms are next specified in turn.

### Toy MDP / Anxiety-to-Depression Transition

Both the Toy MDP and anxiety-to-depression transition simulations were performed using simple gridworlds. Both environments involved only two non-zero states, one rewarding (*r* = 10) and one aversive (*r* = −10). In both environments, we solved for the discounted, asymptomatic *Q*-values using value iteration with *γ* = 0.95. In the toy gridworld, we performed simulations under pessimistic assumptions, *w* ∈ [1.0,0.5,0.0]. In the transition gridworld, we performed simulations under pessimistic assumptions, *w* ∈ [1.0,0.6]. To highlight the effects of learned, we took “snapshots” of *Q*-values prior to asymptote, three and five steps into value iteration.

### Balloon Analog Risk Task / Predator Avoidance Task

The balloon analog risk task^24^ has participants inflate a virtual balloon for points. Earnings rise with each pump, but so too does the risk of the balloon popping and subsequent point loss. Unbeknownst to participants, the number of pumps before balloon pop is predefined and drawn randomly from some distribution (e.g. uniform, normal, exponential), where the mean controls the risk (i.e. average number of pumps before point loss).

Here, we modeled the BART as an undiscounted MDP with 20 states, where transitioning to each successive state yielded *r* =1. The only available actions were to transition to the next state (e.g. *S*_1_ → *S*_2_, *S*_2_ → *S*_3_) or end the episode (i.e. cash-out). With each act to move to the next state, there was some probability of transitioning to a bad terminal state (i.e. balloon pop) with reward equal to the negative equivalent of accumulated gain thus far. The probability of this bad transition was modeled using normal density function, with parameters 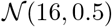 for low risk and 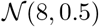 high risk. The asymptomatic *Q*-values were solved for using value iteration for both the low and high risk conditions under pessimistic assumptions, *w* ∈ [1.0,0.6,0.2].

The predator avoidance task^28^ can be analogously modeled. There, a virtual predator approaches participants over discrete time steps while participants “forage”. For every time step the participant remained (i.e. does not flee), points were accumulated. However, if the participant is caught all earnings were lost and an additional penalty was received. Thus, the predator avoidance task bears striking resemblance to the BART; foraging is equivalent to virtual pumps, and fleeing is equivalent to cashing-out.

### Behavioral Inhibition Task

In the behavioral inhibition task^29^ (or sleeping predator task), participants collect virtual tokens while evading capture from a virtual predator. Unlike the BART, the risk in the behavioral inhibition task (i.e. predator “waking up”) is constant. However, the cost of capture increases as sequential tokens are collected.

True to the original, we model the task as an undiscounted MDP with 6 states where transitioning to each successive state yielded *r* =1. Identical to the BART, the only available actions were to transition to the next state or end the episode (i.e. avoid predator). With each act to move to the next state, there was a constant probability of transitioning to a bad terminal state (i.e. capture by predator) with reward equal to the negative equivalent of accumulated gain thus far. The probability of bad transition was defined as *p* = 0.10 for low risk and *p* = 0.15 high risk, based on the objective risk probabilities in the empirical experiment^29^. The asymptomatic *Q*-values were solved for using value iteration for both the low and high risk conditions under pessimistic assumptions, *w* ∈ [1.0,0.6,0.2].

### Aversive Pruning

In the aversive pruning task^32,33^, participants learn to navigate a six-state graphworld, where each state is directly connected to only two other states. Each state is associated with some reward (cost), and participants must plan trajectories through the state-space so as to maximize reward (minimize cost).

We model the task as an undiscounted MDP where the original 6-node network has been “unravelled” into a 15-state decision tree. This is equivalent to having the participant start in one state and plan to make three actions. The rewards associated with transitioning to each state were taken directly from^32,33^. The asymptomatic *Q*-values were solved for using value iteration for both the low and high risk conditions under pessimistic assumptions, *w* ∈ [1.0,0.5].

### Free Choice Task

In the free choice task^44^, participants complete a series of two-stage trials. In the first stage, they select between a free choice option, allowing them to make an additional choice in the second stage, and a fixed choice, permitting no choice in the second stage. In the second stage, participants select between one of two bandits (free choice) or are randomly assigned a bandit (fixed choice). Importantly, all bandits pay out under an identical reward distribution.

We model the task as an undiscounted MDP with 6 state decision tree structure. In the free choice branch, agents are able to select between two terminal bandits; in the fixed choice branch, agents can only choose one terminal bandit. All bandits pay out identically, in this case randomly in the set, *r* ∈ [−1, 1]. Learned *Q*-values were computed using pessimistic temporal difference learning algorithm, with learning rate *η* = 0.4. Each simulated agent learned the values of each action over 100 trials, with an increasing, logarithmically spaced inverse temperature in the range of *β* ∈ [0, 15]. (Inverse temperature was gradually increased over learning to facilitate exploration of choice options.) The resulting fraction of free choices made over the last 50 trials were stored for 1000 simulated agents, run separately for pessimistic assumption, *w* ∈ [1.0,0.5,0.0].

## Acknowledgements

The authors are grateful to Zeb Kurth-Nelson and Will Dabney for helpful discussion. This work was funded in part by NIDA 1R01DA038891, part of the CRCNS program, and by an NSF Graduate Research Fellowship.

## Notes

https://github.com/ndawlab/seqanx

